# Beat-based and memory-based temporal expectations in rhythm: similar perceptual effects, different underlying mechanisms

**DOI:** 10.1101/613398

**Authors:** Fleur L. Bouwer, Henkjan Honing, Heleen A. Slagter

## Abstract

Predicting the timing of incoming information allows the brain to optimize information processing in dynamic environments. Behaviorally, temporal expectations have been shown to facilitate processing of events at expected time points, such as sounds that coincide with the beat in musical rhythm. Yet, temporal expectations can develop based on different forms of structure in the environment, not just the regularity afforded by a musical beat. Little is still known about how different types of temporal expectations are neurally implemented and affect performance. Here, we orthogonally manipulated the periodicity and predictability of rhythmic sequences to examine the mechanisms underlying beat-based and memory-based temporal expectations, respectively. Behaviorally and using EEG, we looked at the effects of beat-based and memory-based expectations on auditory processing when rhythms were task relevant or task irrelevant. At expected time points, both beat-based and memory-based expectations facilitated target detection and led to attenuation of P1 and N1 responses, even when expectations were task-irrelevant (unattended). For beat-based expectations, we additionally found reduced target detection and enhanced N1 responses for events at unexpected time points (e.g., off-beat), regardless of the presence of memory-based expectations or task relevance. This latter finding supports the notion that periodicity selectively induces rhythmic fluctuations in neural excitability and furthermore indicates that while beat-based and memory-based expectations may similarly affect auditory processing of expected events, their underlying neural mechanisms may be different.

---

To optimize sensory processing and perception in a changing environment, the human brain continuously tries to predict incoming information (Clark, 2013; Friston, 2005). Being able to not only predict the content of sensory input (“what”), but also its timing (“when”) allows the system to prepare for and focus on time points when useful information is likely to occur (Large & Jones, 1999; Nobre & van Ede, 2018). Indeed, temporal expectations have been shown to improve processing of events at expected time points (Haegens & Zion Golumbic, 2018; Henry & Obleser, 2012; Nobre & van Ede, 2018; Rohenkohl, Cravo, Wyart, & Nobre, 2012; ten Oever, Schroeder, Poeppel, van Atteveldt, & Zion-Golumbic, 2014). Additionally, temporal expectations allow us to align our actions to sensory input, enabling complex behaviors such as dancing and synchronizing to musical rhythm (Honing & Bouwer, 2019; McGarry, Sternin, & Grahn, 2019; Merchant, Grahn, Trainor, Rohrmeier, & Fitch, 2015).

Temporal expectations are often studied in the context of some form of periodic input, such as a regular beat in music (*beat-based expectations*). However, temporal expectations can be formed based on different types of structure in the environment, which need not necessarily be periodic. For example, temporal expectations can result from learning the relationship between a cue and a particular temporal interval, or from learning (nonperiodic) sequences of temporal intervals (Nobre & van Ede, 2018). In the latter two cases, expectations rely on memory of absolute durations. We will refer to these as *memory-based expectations*. Note that elsewhere, the terms duration-based timing, absolute timing, and interval-based timing have also been used (Breska & Ivry, 2018; Merchant & Honing, 2014; Teki, Grube, Kumar, & Griffiths, 2011).

Temporal expectations have been explained by entrainment models, such as Dynamic Attending Theory (DAT). Such models propose that temporal expectations result from synchronization between internal oscillations and external rhythmic stimulation (Haegens & Zion Golumbic, 2018; Henry & Herrmann, 2014; Jones & Boltz, 1989; Large & Jones, 1999). On a neural level, the internal oscillations can be thought of as fluctuations in low-frequency oscillatory activity, or cortical excitability, such that the high-excitability phase of low frequency neural oscillations coincides with the timing of expected events, facilitating their processing by increasing sensory gain (Lakatos, Karmos, Mehta, Ulbert, & Schroeder, 2008; Schroeder & Lakatos, 2009).

It is unclear whether entrainment, thought to underlie beat-based expectations, can account for memory-based expectations, which do not rely on periodic input (Breska & Deouell, 2017b; Morillon, Schroeder, Wyart, & Arnal, 2016; Rimmele, Morillon, Poeppel, & Arnal, 2018). Arguably, memory-based expectations can better be explained within a predictive processing framework. Notably, beat-based expectations, while often thought to be dependent on entrainment, have also been modeled using hierarchical predictive coding models (Forth, Agres, Purver, & Wiggins, 2016; van der Weij, Pearce, & Honing, 2017). In such models, temporal expectations reflect the learned probability of an event at a given time, similar to learning the probability of content (“what”; Koelsch, Vuust, & Friston, 2019). Thus, whether beat-based and memory-based temporal expectations are based on shared or separate underlying mechanisms – be it entrainment, leading to changes in sensory gain, or predictive processing based on learned probabilistic information – is presently a matter of active debate. In the current EEG study, we addressed this outstanding question by directly comparing the effects of beat-based and memory-based expectations on auditory processing and behavior.

Previously, beat-based and memory-based timing have been differentiated in terms of their occurrence in different species (Honing, Bouwer, Prado, & Merchant, 2018; Honing & Merchant, 2014), the neural networks involved (Teki et al., 2011), and in how they are affected by neuropsychological disorders (Breska & Ivry, 2018), suggesting that beat-based and memory-based expectations are subserved by separate mechanisms. However, some have argued for one integrated system for timing (Schwartze & Kotz, 2013; Teki, Grube, & Griffiths, 2012), based on neuropsychological evidence (Cope, Grube, Singh, Burn, & Griffiths, 2014), and for reasons of parsimony (Rimmele et al., 2018). Complicating this discussion, to date, many studies examining temporal expectations used isochronous sequences of events to elicit temporal expectations in general, and beat-based expectations in particular (Arnal, Doelling, & Poeppel, 2014; Breska & Deouell, 2016; Breska & Ivry, 2018; Henry & Obleser, 2012; Lakatos et al., 2013; Lawrance, Harper, Cooke, & Schnupp, 2014; Rohenkohl et al., 2012; Schwartze, Farrugia, & Kotz, 2013; Teki et al., 2011). Isochronous sequences are fully predictable, both in terms of their absolute intervals, and in terms of their ongoing periodicity, and therefore do not allow for differentiation between beat-based and memory-based expectations.

A handful of studies attempted to directly compare beat-based and memory-based expectations by using both isochronous stimuli, and stimuli that were predictable, but not periodic. In a behavioral experiment, responses to sequences that were isochronous (affording both beat-based and memory-based expectations), predictably speeding up or slowing down (affording only memory-based expectations), or with random timing (no expectations) were compared (Morillon et al., 2016). Beat-based expectations improved both perceptual sensitivity and response speed, while memory-based expectations only affected perceptual sensitivity, which was suggested to result from a special relationship between beat-based expectations and the motor system (Morillon et al., 2016). However, in another behavioral study, both beat-based and memory-based expectations improved response speed (Breska & Ivry, 2018), and phase coherence of delta oscillations, which is often used as a proxy for neural entrainment, was shown to be similarly enhanced by memory-based and beat-based expectations (Breska & Deouell, 2017b), suggesting either that phase coherence is not a good measure of entrainment, as suggested by the authors, or that entrainment is a general, rather than a context-specific, mechanism of temporal expectations (Rimmele et al., 2018). Note that in all three studies described above (Breska & Deouell, 2017b; Breska & Ivry, 2018; Morillon et al., 2016), while care was taken to design stimuli that elicited only memory-based but not beat-based expectations, responses to isochronous stimuli were used as a proxy for beat-based expectations. In addition to affording beat-based expectations, isochronous stimuli are by definition more predictable in terms of learning their intervals than any other type of memory-based sequence (e.g., only one interval needs to be learned in an isochronous sequence). Thus, in these studies, the effects of beat-based expectations were always confounded with increases in temporal predictability based on memory. In addition, in the latter two studies (Breska & Deouell, 2017b; Breska & Ivry, 2018), static visual stimuli were used. The auditory system has repeatedly been found to be superior over the visual system in eliciting temporal expectations in humans (Grahn, 2012; Grahn, Henry, & McAuley, 2011; Zarco, Merchant, Prado, & Mendez, 2009). While it is possible to achieve equal synchronization performance (a measure of temporal expectations) with visual and auditory stimuli alike, this requires the use of moving, rather than static visual input (Grahn, 2012; Iversen, Patel, Nicodemus, & Emmorey, 2015). Thus, the stimuli used by previous studies to study beat-based expectations not only allowed for memory-based strategies to form expectations, they were arguably not optimal for creating beat-based expectations at all.

Here, to gain a better understanding of how beat-based and memory-based expectations may (differentially) influence early sensory processing, we orthogonally manipulated beat-based and memory-based expectations using auditory stimuli, and examined their effects on both behavioral and auditory ERP responses. Effects on ERPs have not been studied before in this context and may provide a potentially fruitful way of differentiating between beat-based and memory-based expectations. Temporal expectations have been reported to both enhance and attenuate the auditory P1 and N1 responses. Enhancement of sensory responses at expected time points (Bouwer & Honing, 2015; Escoffier et al., 2015; Fitzroy & Sanders, 2015; Hsu, Hämäläinen, & Waszak, 2013; Rimmele, Jolsvai, & Sussman, 2011; Tierney & Kraus, 2013) is in line with entrainment models of temporal expectations, which assume increased sensory gain at expected time points (Large & Jones, 1999). By contrast, attenuation of sensory responses at expected time points (Lange, 2009; Paris, Kim, & David, 2016; Sanabria & Correa, 2013; Schwartze et al., 2013; Sherwell, Garrido, & Cunnington, 2017; van Atteveldt et al., 2015) is in line with predictive models of brain function that assert more efficient processing of incoming information when predicted information is suppressed (Friston, 2005; Marzecová, Widmann, Sanmiguel, Kotz, & Schröger, 2017; Schröger, Kotz, & SanMiguel, 2015; Schröger, Marzecová, & Sanmiguel, 2015). Whether temporal expectations lead to enhancement or attenuation of sensory responses may depend on the type of temporal structure that affords an expectation. Periodic input, affording beat-based expectations, may lead to entrainment and thus increased neural excitability at expected time points, resulting in enhancement of early sensory responses (Haegens & Zion Golumbic, 2018). By contrast, learned probabilistic information about timing, affording memory-based expectations, may lead to the suppression of predicted information, resulting in attenuation of early sensory responses. Thus, if based on separate mechanisms, one subserved by entrainment and one by predictive processing, beat-based and memory-based expectations may have opposing effects on sensory responses, enhancement and attenuation of responses respectively.

In addition to the effects of beat-based and memory-based expectations on behavioral and auditory responses, we examined the effects of task relevance on both types of expectations. Entrainment and beat-based processing have been shown to be somewhat independent of task relevance (Bouwer, Van Zuijen, & Honing, 2014; Bouwer, Werner, Knetemann, & Honing, 2016; Breska & Deouell, 2014; Rohenkohl, Coull, & Nobre, 2011). Predictive processing, however, has been shown to depend on and interact with task relevance in both the visual and the auditory domain (Hsu, Hämäläinen, & Waszak, 2018; Kok, Rahnev, Jehee, Lau, & de Lange, 2012; Paris et al., 2016; Todorovic, Schoffelen, Ede, Maris, & de Lange, 2015). Thus, if beat-based expectations rely on entrainment while memory-based expectations rely on learning probabilistic information, they may be affected by task relevance differently.

In the current study, in two experiments, we compared responses to auditory sequences that were either periodic, affording beat-based expectations, or aperiodic, thus not affording beat-based expectations. Also, sequences could either consist of fully predictable temporal intervals, affording memory-based expectations, or unpredictable, randomly concatenated intervals. The responses to events in the rhythmic sequences thus depended on their expectedness, with expectations coming from two distinct sources: the periodicity of the sequence (beat-based) and the repetition of the pattern (memory-based). We not only examined responses for events on the beat (e.g., in phase with the periodicity, at expected times), but also off the beat (e.g., out of phase with the periodicity, at less expected times), as entrainment theories predict not only increased sensory gain at expected moments, but also reduced sensory gain in between (Breska & Deouell, 2014). Thus, on the beat, when comparing periodic with aperiodic sequences, we could assess the facilitating effects of the presence of beat-based expectations, while off the beat, we could assess the effects of events occurring at times that were “mispredicted” in terms of the expected beat (Hsu, Bars, & Ha, 2015).

In Experiment 1, we examined the effects of beat-based and memory-based expectations at two positions (on and off the beat) on behavioral performance, by measuring the speed and accuracy of the detection of targets in the form of rare softer tones in the rhythmic sequences. In Experiment 2, we additionally recorded ERP responses to all non-target sounds, to examine the effects of beat-based and memory-based expectations at two positions on P1 and N1 responses. Additionally, in Experiment 2, ERP responses were examined both when sequences were task-relevant and task-irrelevant.

In terms of behavioral outcomes, we expected faster and more accurate detection of targets with expected than unexpected timing, both for beat-based and memory-based expectations. Additionally, beat-based and memory-based expectations may interact in two distinct ways. First, the presence of beat-based expectations may lead to an increase in sensory gain for sounds on the beat, as predicted by entrainment models. Increased gain should increase the precision of memory-based predictions (Feldman & Friston, 2010), leading to enhanced effects of memory-based expectations in the presence of beat-based expectations. Thus, interacting effects of beat-based and memory-based expectations, which are larger than could be expected based on additivity, could indicate separate mechanisms, with beat-based expectations influencing the gain or precision of sensory processing through entrainment, and memory-based expectations affecting the probabilistic predictions themselves. Alternatively, if beat-based and memory-based expectations rely on shared mechanisms, the simultaneous presence of both types of expectations may lead to interference, with smaller effects of either type when both need to be engaged. In terms of ERP responses, qualitative differences between the effects of beat-based and memory-based expectations – the former leading to enhancement, and the latter to attenuation of sensory responses – could indicate different underlying mechanisms. However, if both types of expectations rely on similar mechanisms, be it based on entrainment or predictive processing, we expected them to affect sensory responses similarly.

## Methods

### Experiment 1

#### Participants

Thirty-four participants (26 women), aged between 19 and 45 years old (M = 24.6, SD = 5.7) with no history of neurological or hearing disorders took part in the experiment. Data from two participants were removed due to technical problems, leaving 32 participants for the analysis. All participants provided written consent prior to the study, and participants were reimbursed with either a monetary fee or course credit. The study was approved by the Ethics Review Board of the Faculty of Social and Behavioral Sciences of the University of Amsterdam. The statistical analysis of Experiment 1 was preregistered (https://aspredicted.org/az8kr.pdf).

#### Stimuli

We created sound patterns of five or six consecutive temporal intervals (Figure 1A), marked by identical woodblock sounds of 60 ms length, generated in GarageBand (Apple Inc.). Patterns of five or six intervals are short enough to allow for learning of the temporal intervals (Schultz, Stevens, Keller, & Tillmann, 2013), and to not make too large demands on working memory (Grahn & Schuit, 2012). At the same time, with a total length of 1800 ms, patterns were long enough to avoid the perception of a regular, periodic beat when patterns were concatenated into sequences, as people do not readily perceive a beat with a period of 1800 ms (London, 2012). Patterns were concatenated into sequences of 128 patterns, with a final tone added to each sequence. Each sequence thus lasted for 3 minutes and 51 seconds.

**Figure 1.**
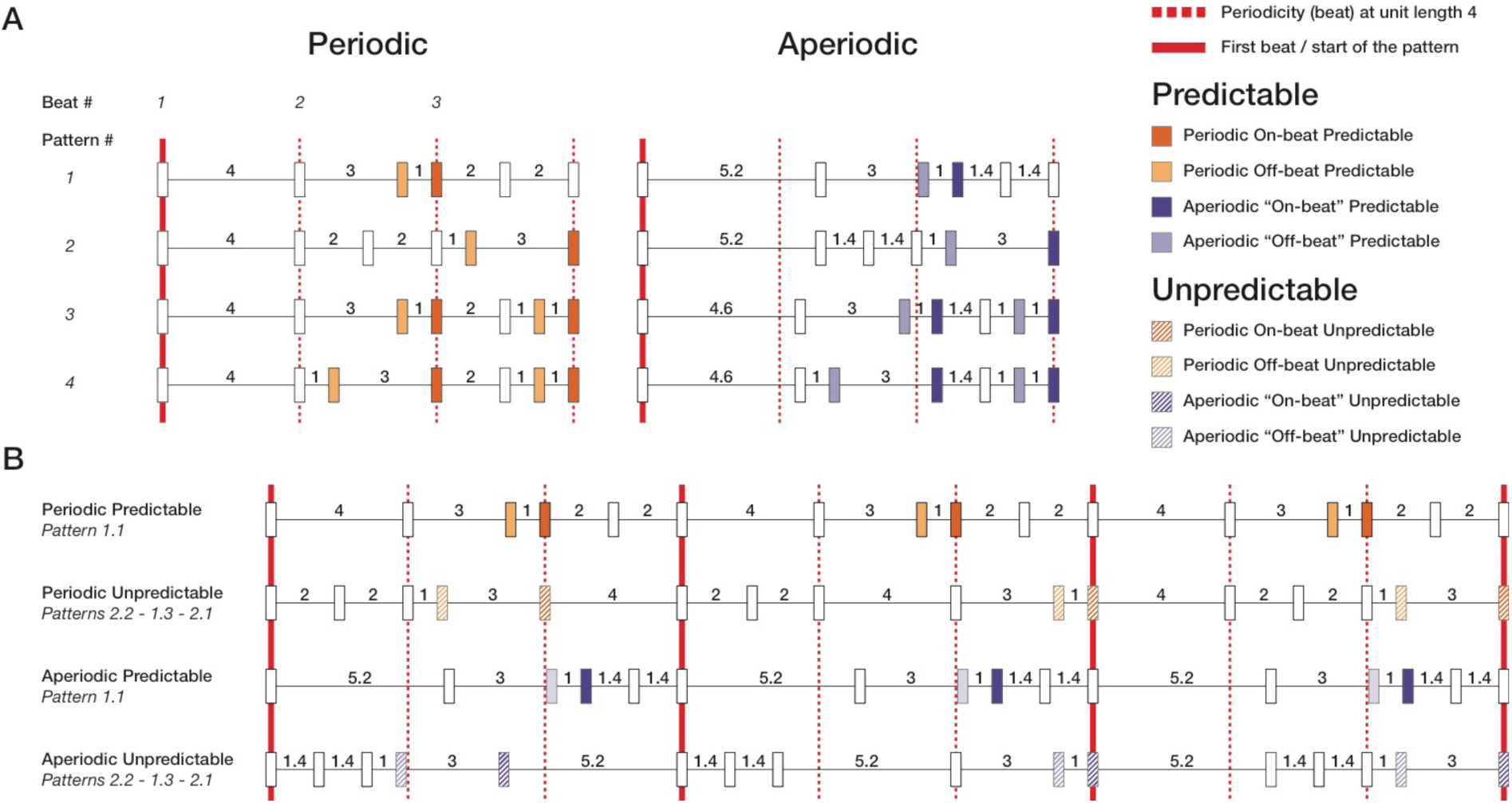
Schematic overview of the stimuli. A) Patterns of 5 (patterns 1 and 2) and 6 (patterns 3 and 4) temporal intervals were created. All patterns were 12 units length, equaling 1800 ms (1 unit = 150 ms). Periodic patterns (left, orange shades) consisted of intervals related by integer ratios 1:2:3:4. In periodic patterns, groups of intervals always added up to unit length 4, creating a beat with an inter-beat onset of 600 ms (red dotted lines). Aperiodic patterns (right, purple shades) consisted of intervals related by non-integer ratios, not allowing for the perception of a regular beat. B) Patterns were concatenated into sequences of either 128 identical patterns (predictable sequences), or 128 randomly chosen patterns (unpredictable sequences). For the unpredictable sequences, not only patterns starting at beat 1 (patterns 1.1, 2.1, 3.1, and 4.1) were used, but also cyclic permutations of these patterns, starting at beats 2 or 3 (patterns 1.2, 1.3, 2.2, etc.). For all analyses, only events after intervals with unit lengths 1 and 3 were used, to control for acoustic context.

##### Beat-based expectations

For the periodic, beat-based patterns, temporal intervals were related by the integer ratios of 1:2:2:3:4 (five intervals) and 1:1:1:2:3:4 (six intervals). The shortest interval was set at 150 ms, leading to inter-onset intervals for the other intervals of 300, 450, and 600 ms. In the periodic patterns, temporal intervals were organized to form groups of four units length (600 ms) and grouped such that a perceptually accented tone was present at the start of each group (Grahn & Brett, 2007; Povel & Essens, 1985). In these patterns a beat could be perceived with an inter-beat interval of 600 ms (100 BPM or 1.7 Hz), the optimal rate for human beat perception (London, 2012). These patterns could be regarded as strictly metric, with the periodicity of the pattern always being marked by a sound (Grahn & Brett, 2007). Each pattern consisted of 12 units length, or three beats of four units length.

To create aperiodic equivalents of the beat-based patterns, we changed the ratios by which the temporal intervals were related. For the aperiodic patterns, intervals were related by non-integer ratios of 1:1.4:1.4:3:5.2 (five intervals) and 1:1:1:1.4:3:4.6 (six intervals). The aperiodic patterns were equal to their periodic counterparts in terms of length, grouping, and number of tones. However, the aperiodic patterns did not contain a periodic beat at unit length four. A pilot confirmed that aperiodic patterns were rated as less beat-inducing than periodic patterns.

Note that it is impossible to create sequences of sounds that are not to some extent (quasi-) periodic (Breska & Deouell, 2017a; Obleser, Henry, & Lakatos, 2017). However, the sequences clearly differed in the presence of periodicity at a rate that afforded beat-based expectations. First, in the aperiodic patterns, contrary to the periodic patterns, events did not align with the intended beat at a rate close to the ideal tempo for beat perception (1.6 Hz). This was also apparent from a spectral analysis of the waveforms (results not shown). Peaks at 1.6 Hz (the beat frequency) were larger for the beat-based sequences than the memory-based sequences. Second, while periodicity was present in the aperiodic sequences at the level of concatenated patterns (with a period of 1800 ms, or 33 BPM or 0.6 Hz), this was too slow for humans to readily perceive a beat in (London, 2012). Finally, we performed an informal pilot experiment in which we asked 17 participants to rate on a scale from 1 to 10 how strongly they heard a beat in the aperiodic and periodic patterns. On average, each periodic pattern was rated as containing more beat (on average 9.3) than each aperiodic pattern (on average 6.7), confirming that our manipulation of periodicity indeed affected perception at the level of hearing a beat. Thus, while we are aware that the aperiodic patterns could be classified as quasi or weakly periodic, for clarity, we will refer to them as aperiodic.

##### Memory-based expectations

Fully predictable sequences were created by concatenating 128 identical patterns into a sequence (Figure 1B). The surface structure of temporal intervals in these sequences could easily be predicted based on probabilistic information alone. Unpredictable sequences were created by concatenating 128 semi-randomly chosen patterns. Patterns were chosen both from the original patterns, which were also used for the predictable sequences (patterns starting at beat 1, see Figure 1), and from cyclic permutations of these (patterns starting at beats 2 or 3, see Figure 1). The cyclic permutations were identical to the original patterns when looped (as in the predictable sequences), but not when concatenated in random order (as in the unpredictable sequences). Within an unpredictable sequence, only patterns with either five or six intervals were concatenated. The longest interval had a different length in the aperiodic patterns with five and six intervals (unit lengths 5.2 and 4.6 respectively). This was necessary to keep the overall length of each pattern identical. However, combining the two sets would lead to a larger number of possible temporal intervals in the aperiodic than periodic sequences, possibly confounding the effects of beat-based expectations with differences in entropy. Thus, to keep both the number of events per pattern (event density) and the number of possible intervals (entropy) identical between conditions, we did not combine the sets of rhythms with five and six intervals. Within unpredictable sequences, each pattern could occur maximally twice consecutively.

##### Position

With manipulations of periodicity and predictability, we were able to compare responses to events that were expected based on a beat or on learned interval structure with responses that could not be predicted based on their timing. However, we also wanted to examine how beat-based expectations affected responses to events with unexpected timing (e.g., not unpredicted, but rather mispredicted, see also Hsu et al., 2018). Therefore, we not only probed events that were in phase with the periodicity (e.g., on the beat), but also events that were out of phase with the periodicity (e.g., off the beat, see Figure 1). The off-beat events fell at quarter-phase (150 or 450 ms) relative to the beat. We did not include events at antiphase (300 ms), as possibly, participants could perceive an additional beat at the faster subdivision rate, making these events ambiguous in terms of their metrical salience.

We assumed that people would not perceive a beat in the aperiodic patterns, and therefore, that the distinction between on the beat and off the beat would be meaningless for these patterns in terms of temporal expectations. However, grouping effects could lead to differences in the perception of events on and off the beat. For example, the auditory system tends to group events in groups of two (Abecasis, Brochard, Granot, & Drake, 2005; Brochard, Abecasis, Potter, Ragot, & Drake, 2003; Potter, Fenwick, Abecasis, & Brochard, 2009), and perceptual accents are known to arise based on the surface structure of a rhythm (Povel & Essens, 1985). To be able to assess the effects of temporal expectations at different positions without confounding these with differences in grouping, we classified events in the aperiodic patterns as on-beat or off-beat, depending on their grouping in the periodic counterpart, even when we assumed no beat would be perceived in response to the aperiodic patterns (i.e., when we refer to an event as “on-beat” in an aperiodic pattern, it is an event that falls on the beat in the periodic equivalent, and thus is matched to its periodic equivalent in terms of grouping).

##### Targets

In the behavioral task, temporal expectations were probed implicitly, by introducing infrequent intensity decrements as targets. Based on previous experiments, we expected that temporal expectations would improve the detection of these targets (Bouwer & Honing, 2015; Bouwer et al., 2014, 2016; Potter et al., 2009). Intensity decrements of 6 dB were used (Bouwer & Honing, 2015). In each sequence of 128 patterns, 32 patterns (25 percent) contained a target. Half of the targets appeared on the beat, and half of the targets off the beat. In each sequence, 26 targets were in positions after temporal intervals with unit lengths 1 and 3, present in both periodic and aperiodic patterns. Only these targets were used for the analysis, to equate their acoustic context. At least two standard patterns separated a pattern containing a target.

##### Procedure

A total of 16 sequences were presented to each participant, four of each type. Sequences of different types were semi-randomized, with each type appearing once every four sequences, and therefore a maximum of two sequences of the same type in a row. Upon arrival, participants completed a consent form and were allowed to practice the task. They were instructed to avoid movement, listen to the rhythm carefully, and press a button as fast as possible when they heard a target. Participants were allowed breaks between sequences. An entire experimental session lasted for about 2 hours. Participants were tested individually in a dedicated lab at the University of Amsterdam. Sounds were presented at 70 dB SPL with one Logitech speaker positioned in front of the participants, using Presentation software (version 19.0, www.neurobs.com).

#### Data analysis

In total, each participant was presented with 52 targets for each condition that were included in the analysis. All responses made within 2000 ms of a target were recorded. Responses faster than 150 ms were discarded, as were responses that were more than 2.5 standard deviations from the mean of a participant’s reaction time within each condition. Removal of outliers led to the exclusion of 2.9 percent of the responses in Experiment 1 and 3.1 percent of the responses in Experiment 2. Participants who did not achieve a hit rate of higher than 50 percent in any of the conditions were excluded from the analysis. In Experiment 1, on this ground, two participants were excluded, leaving 30 participants for the analysis of hit rates. One additional participant was excluded for the analysis of reaction times, as this participant had less than five valid reaction times in one condition (i.e., less than 5 out of a possible 52 targets were hits). In Experiment 2, no participants were excluded from the analysis. Hit rates for each condition and participant and average reaction times for each condition and participant were entered into three-way repeated measures ANOVAs, with Periodicity (periodic, aperiodic), Predictability (predictable, unpredictable), and Position (on the beat, off the beat) as within-subject factors.

The meaning of the factor Position can be regarded as somewhat ambiguous. In the beat-based sequences, differences between on-beat and off-beat positions could be due to both differences in grouping, and differences in metrical position (assuming that participants would hear a beat in these sequences). In the memory-based sequences, grouping differences could still affect the results for the factor Position, while there would presumably not be an effect of metrical position. Thus, the meaning of the factor Position depends on the Periodicity of a sequence, potentially limiting the conclusions that can be drawn from an ANOVA that assumes this factor to be orthogonal to the other factors of interest. Therefore, in addition to the (pre-registered) full ANOVA, we also ran separate ANOVAs for on-beat and off-beat positions, to check whether our results would hold up when taking into account the possible confounded Position factor. For all ANOVAs, for significant interactions (p<0.05), post hoc tests of simple effects were performed. For all simple effects, we report uncorrected p-values. Effect sizes are reported as partial eta squared. All statistical analyses were performed in SPSS 24.

### Experiment 2

#### Participants

Thirty-two participants (22 women), aged between 19 and 44 years old (M = 23.4, SD = 4.9) with no history of neurological or hearing disorders took part in Experiment 2. Data of one participant were removed due to excess noise in the EEG signal, leaving thirty-one participants for the analysis. All participants provided written consent prior to the study, and participants were reimbursed with either a monetary fee or course credit. The study was approved by the Ethics Review Board of the Faculty of Social and Behavioral Sciences of the University of Amsterdam.

#### Stimuli and Procedure

The materials and procedure for Experiment 2 were identical to those for Experiment 1. In Experiment 2, participants additionally completed an unattended version of the experiment. In the unattended condition, participants were asked to ignore the rhythmic sequences and focus on a self-selected muted movie, rendering the stimuli task irrelevant. All participants first completed the unattended EEG experiment, and subsequently the attended EEG experiment. The EEG recording lasted for around 2 hours. Participants were encouraged to take breaks if needed. During the recording, data quality was assessed online by the experimenter, and if needed, channels were refitted and extra electrode gel was applied. For Experiment 2, one experimental session was about 4 hours, including breaks, practice, and setting up equipment.

#### EEG recording

EEG was recorded using a 64-channel Biosemi Active-Two acquisition system (Biosemi, Amsterdam, The Netherlands), with a standard 10/20 configuration and additional electrodes for EOG channels, on the nose, on both mastoids, and on both earlobes. The EEG signal was recorded at 1 kHz.

#### EEG analysis

Preprocessing was performed in MATLAB and EEGLAB (Delorme & Makeig, 2004). Data were offline re-referenced to linked mastoids, bad channels were removed, and independent component analysis was used to remove eye-blinks. Subsequently, bad channels were replaced by values interpolated from the surrounding channels. Visual inspection of the ERPs revealed a postauricular muscle response (PAM) in several subjects. The auditory evoked potential can be easily contaminated by the PAM response (Bell, Smith, Allen, & Lutman, 2004; Picton, Hillyard, Krausz, & Galambos, 1974). To avoid contamination, we re-referenced the data to earlobes for all further analyses. For completeness, we also report results from the mastoid-referenced data.

Data were offline down-sampled to 512 Hz, and filtered using 0.1 Hz high-pass and 40 Hz low-pass finite impulse response filters. Epochs for each condition separately were extracted for non-target sounds, from 200 ms preceding the onset of each event till 500 ms after the onset of each event. Only epochs for events following an interval of unit length 1 or unit length 3 (150 and 450 ms respectively) were included, to equate the acoustic context of events used in the analysis. Also, the first 12 sounds of each sequence (2 whole patterns) were excluded from the analysis, to allow for the build-up of expectations. Epochs with a voltage change of more than 150 microvolt in a 200 ms sliding window were rejected from further analysis. For each condition and participant, epochs were averaged to obtain ERPs and baseline corrected using the average voltage of the 50 ms window preceding each sound. Finally, ERPs were averaged over participants to obtain grand average waveforms.

Peak latencies for the P1 and N1 responses were determined independent from the statistical analysis, from the average waveform collapsed over all conditions. P1 peaked at 58 ms after tone onset. We defined P1 amplitude as the average amplitude in a 20 ms window around the peak (48-68 ms). N1 peaked at 124 ms, and was more distributed in time. Thus, we defined N1 amplitude as the average amplitude from a 40 ms window around the peak (104-144 ms). Auditory evoked potentials are known to be maximal over fronto-central electrodes (Picton et al., 1974; Ruhnau, Herrmann, Maess, & Schröger, 2011), which was also observed in the current dataset. Therefore, ERP amplitudes were computed from the average of a cluster of 15 fronto-central electrodes: F3, F1, Fz, F2, F4, FC3, FC1, FCz, FC2, FC4, C3, C1, Cz, C2, and C4. All statistics and figures reported here are based on the average amplitude from this region of interest.

#### Statistical analysis

Amplitudes for P1 and N1 were entered into repeated measures ANOVAs, with four within-subject factors: Periodicity (periodic, aperiodic), Predictability (predictable, unpredictable), Position (on the beat, off the beat), and Attention (attended, unattended). As for the behavioral results, we also included separate ANOVAs for on-beat and off-beat positions, to account for the possible collinearity of Position with Periodicity. For significant interactions (p<0.05), subsequent tests of simple effects were performed. For all simple effects, we report uncorrected p-values. Effect sizes are reported as partial eta squared. Analyses were performed in SPSS 24.

#### Comparing beat-based and memory-based expectations directly

While the factorial design included both Periodicity and Predictability as factors, the ANOVAs do not allow for a direct comparison of their effects. To directly compare the effects of beat-based and memory-based expectations, we performed two additional analyses. In both analyses, the effect of beat-based expectations was quantified as the difference between responses on the beat in periodic and aperiodic sequences. The effect of memory-based expectations was quantified as the difference between responses in predictable and unpredictable sequences. For the latter, we only included responses on the beat, to make sure that possible differences between beat-based and memory-based expectations could not be attributed to differences in grouping.

First, we compared the effects of beat-based and memory-based expectations in the P1 and N1 windows directly using paired-samples T-tests. To quantify the possibility that no differences were present, we performed both traditional and Bayesian T-tests using JASP (JASP Team, 2019; Wagenmakers et al., 2018). We estimated Bayes factors using a Cauchy prior distribution (r = 0.71), with as a null hypothesis no differences between the effects of beat-based and memory-based expectations. In addition, we performed a robustness check as implemented in JASP, to assess whether the results would change with a different prior (r = 1) as proposed originally for Bayesian T-tests (Jeffreys, 1961; Wagenmakers et al., 2018).

Second, we directly compared the effects of memory-based and beat-based expectations using cluster-based permutation tests. With this approach we could examine potential differences at all timepoints and at all electrodes, while taking into account the multiple comparisons along both the spatial and time axes. As ERP components often overlap in time, their real peaks may be obscured in grand average waveforms (Luck, 2005). The use of cluster-based permutation testing allowed us to make sure we did not miss potential differences between beat-based and memory-based expectations by selectively examining peak time windows and selected clusters of electrodes. Cluster-based permutation tests were performed using the Fieldtrip toolbox (Oostenveld, Fries, Maris, & Schoffelen, 2011). All timepoints in the 150 ms time window following the onset of sound events were included. We limited the analysis to the first 150 ms, to avoid contamination of subsequent sounds, which in some but not all conditions, could occur 150 ms following an event. Clusters were formed based on adjacent time-electrode samples that survived a statistical threshold of p<0.01 when comparing the conditions of interest with dependent samples T-tests. Clusters were subsequently evaluated with permutation tests, using 2000 iterations.

## Results

### Behavioral results

Hit rates and reaction times for all conditions from both Experiment 1 and 2 are shown in Table 1 and Table 2 and Figure 2. In Experiment 2, all results from Experiment 1 were replicated with comparable effect sizes.

**Table 1.** Hit rates for all conditions in Experiment 1 and Experiment 2. Hit rates are reported as percentage hits. Standard deviations in brackets. *Simple effect significant at p<0.05 (uncorrected). **Simple effect significant at p<0.01 (uncorrected).

|  |  | Experiment 1 (N=30) |  |  | Experiment 2 (N=31) |  |  |
| --- | --- | --- | --- | --- | --- | --- | --- |
|  |  | Periodic | Aperiodic | Effect of beat-based expectations | Periodic | Aperiodic | Effect of beat-based expectations |
| Beat | Predictable | 82 (14) | 81 (15) | +1 | 79 (17) | 79 (17) | 0 |
|  | Unpredictable | 73 (19) | 69 (23) | +4 | 72 (17) | 67 (20) | +5** |
| Effect of memory-based expectations |  | +9** | +12** |  | +7** | +12** |  |
| Offbeat | Predictable | 63(20) | 68 (21) | -5* | 64 (21) | 72 (19) | -8** |
|  | Unpredictable | 52 (22) | 61 (26) | -9** | 51 (20) | 60 (21) | -9** |
| Effect of memory-based expectations |  | +11** | +7* |  | +13** | +12** |  |

**Table 2.** Reaction Times in ms for all conditions in Experiment 1 and Experiment 2. Standard deviations in brackets. *Simple effect significant at p<0.05 (uncorrected). **Simple effect significant at p<0.01 (uncorrected).

|  |  | Experiment 1 (N=29) |  |  | Experiment 2 (N=31) |  |  |
| --- | --- | --- | --- | --- | --- | --- | --- |
|  |  | Periodic | Aperiodic | Effect of beat-based expectations | Periodic | Aperiodic | Effect of beat-based expectations |
| Beat | Predictable | 538 (78) | 541 (78) | -3 | 517 (59) | 509 (60) | +8 |
|  | Unpredictable | 557 (74) | 571 (80) | -14 | 542 (66) | 554 (75) | -12 |
| Effect of memory-based expectations |  | -19* | -30** |  | -25** | -45** |  |
| Offbeat | Predictable | 573 (84) | 571 (81) | +2 | 550 (93) | 531 (76) | +19 |
|  | Unpredictable | 601 (92) | 591 (96) | +10 | 569 (81) | 571 (96) | -2 |
| Effect of memory-based expectations |  | -28* | -20* |  | -19 | -40** |  |

**Figure 2.**
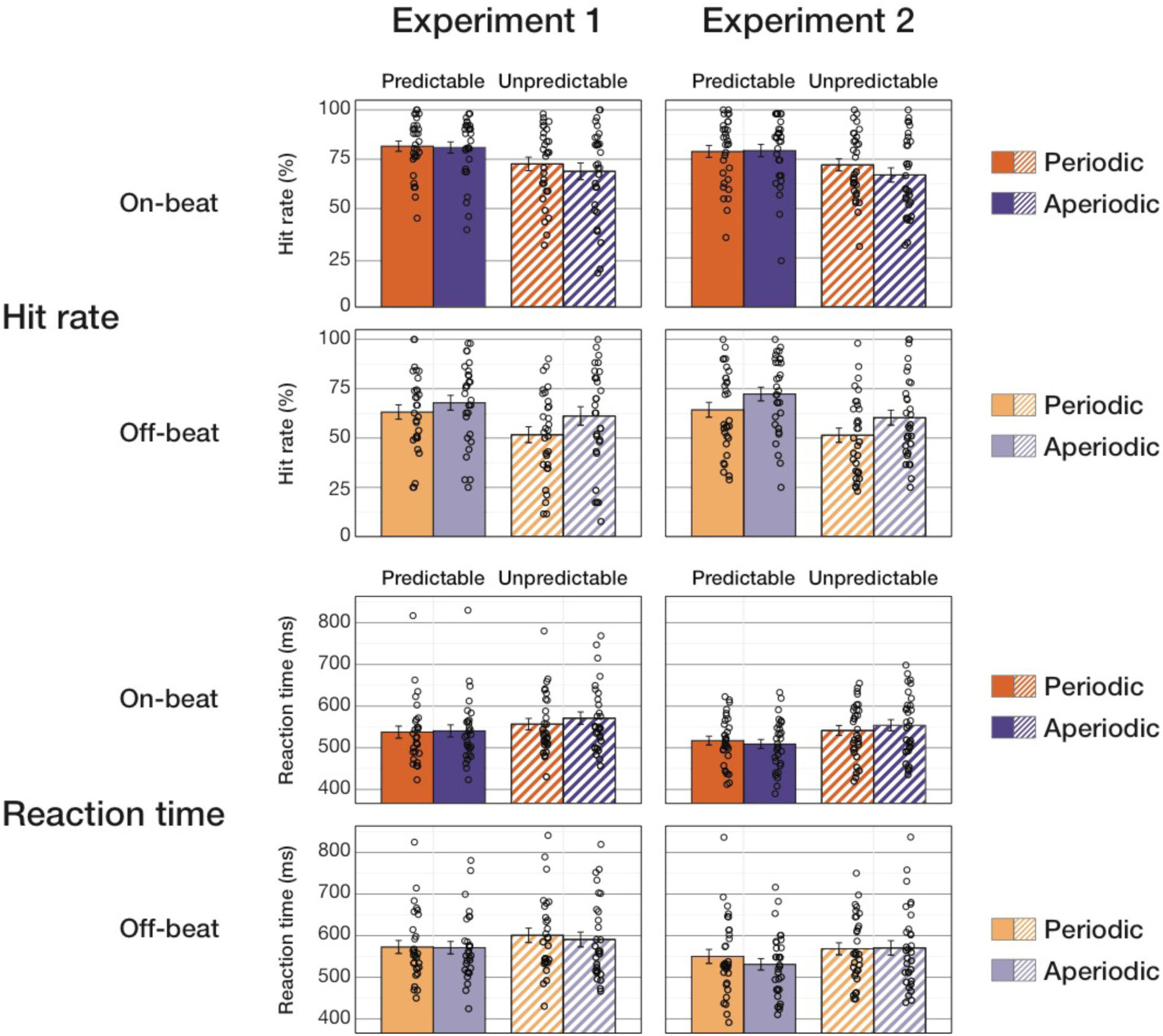
Hit rates (top) and reaction times (bottom) show improved target detection for events at expected time points. Memory-based expectations lead to better and faster target detection in Experiment 1 (left) and Experiment 2 (right). Beat-based expectations only lead to improvements in the absence of memory-based expectations in Experiment 2, but consistently lead to deteriorated detection of targets off the beat in both experiments.

#### Memory-based expectations

In general, targets were detected more often and faster in predictable than in unpredictable sequences, as reflected in a significant main effect of Predictability for hit rates (Experiment 1: *F*(1,29) = 39.4, *p* < 0.001, η_p_^2^ = 0.58; Experiment 2: *F*(1,30) = 46.4, *p* < 0.001, η_p_^2^ = 0.61) and reaction times (Experiment 1: *F*(1,28) = 19.6, *p* < 0.001, η_p_^2^ = 0.41; Experiment 2: *F*(1,30) = 25.0, *p* < 0.001, η_p_^2^ = 0.46). The simple effect of Predictability was significant for all comparisons for hit rates (all uncorrected *p*s < 0.016, see Table 1) and reaction times (all uncorrected *p*s < 0.023, except for targets off the beat in periodic sequences in Experiment 2; *p* = 0.055). Thus, as expected, memory-based expectations led to improved detection of targets at expected time points. Results from the separate ANOVAs for on-beat and off-beat positions confirmed the main effect of Predictability for hit rates (Experiment 1, on-beat: *F*(1,29) = 31.0, *p* < 0.001, η_p_^2^ = 0.52; off-beat: *F*(1,29) = 27.29, *p* < 0.001, η_p_^2^ = 0.49; Experiment 2, on-beat: *F*(1,30) = 35.68, *p* < 0.001, η_p_^2^ = 0.54; off-beat: *F*(1,30) = 34.86, *p* < 0.001, η_p_^2^ = 0.54) and reaction times (Experiment 1, on-beat: *F*(1,28) = 16.6, *p* < 0.001, η_p_^2^ = 0.37; off-beat: *F*(1,28) = 12.4, *p* = 0.001, η_p_^2^ = 0.31; Experiment 2, on-beat: *F*(1,30) = 39.44, *p* < 0.001, η_p_^2^ = 0.57; off-beat: *F*(1,30) = 10.27, *p* = 0.003, η_p_^2^ = 0.26). Figure 3 (top) shows the main effect of Predictability on target detection performance collapsed over the levels of Periodicity and Position.

**Figure 3.**
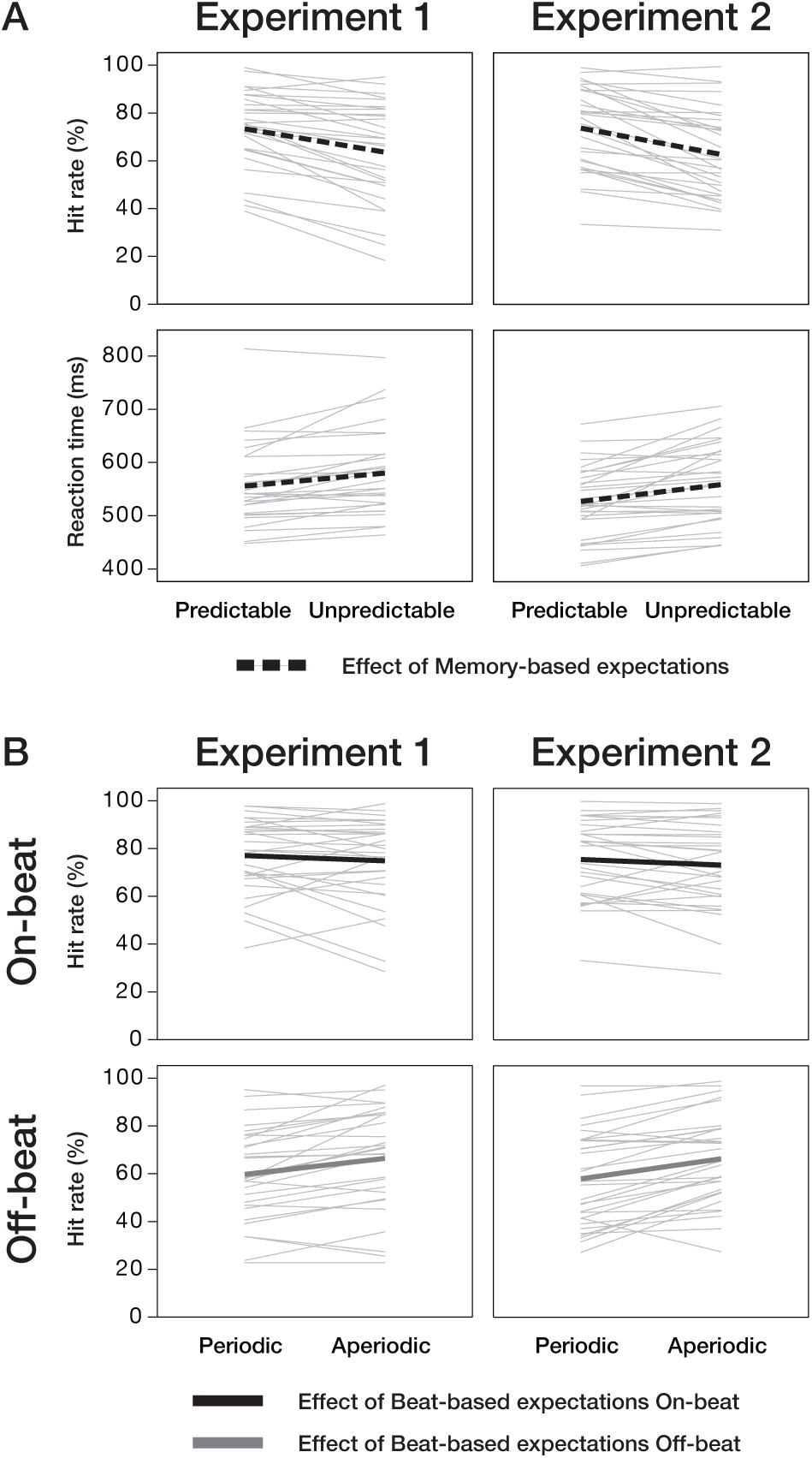
Temporal expectations improve target detection. Memory-based expectations (A) led to both higher hit rates and faster reaction times (main effect of Predictability). Beat-based expectations mainly affected hit rates by deteriorating detection for mispredicted events (off-beat; bottom of panel B). Thick lines represent the mean, thin lines are single participants.

#### Beat-based expectations

In Experiment 2, we found a main effect of Periodicity on hit rates (*F*(1,30) = 6.59, *p* = 0.016, η_p_^2^ = 0.18). However, as expected, the effect of Periodicity on hit rates in both experiments depended on the position of the target. On the beat, targets were detected more often in periodic than aperiodic sequences, while off the beat, targets were detected more often in aperiodic than periodic sequences, as reflected in a significant interaction in the overall ANOVA between Periodicity and Position (Experiment 1: *F*(1,29) = 35.0, *p* < 0.001, η_p_^2^ = 0.54; Experiment 2: *F*(1,30) = 39.3, *p* < 0.001, η_p_^2^ = 0.57). In line with the results for hit rates, numerically, detection of targets off the beat, but not on the beat, was slower in periodic than aperiodic sequences. However, the interaction between Periodicity and Position did not reach significance for the reaction times (Experiment 1: *F*(1,28) = 3.74, *p* = 0.063, η_p_^2^ = 0.12; Experiment 2: *F*(1,30) = 1.91, *p* = 0.18, η_p_^2^ = 0.06). The separate ANOVAs for on-beat and off-beat positions showed that while the facilitating effect of Periodicity on the beat did not reach significance for hit rates, nor for reaction times in either experiment (all *p*s > 0.11), the decrease in performance for targets presented out of phase with the periodicity (e.g., mispredicted) was significant for hit rates, apparent from a main effect of Periodicity for off-beat targets (Experiment 1: *F*(1,29) = 19.71, *p* < 0.001, η_p_^2^ = 0.41; Experiment 2: *F*(1,30) = 30.28, *p* < 0.001, η ^2^ = 0.50). Figure 3 (bottom) shows the main effect of Periodicity on target detection performance separate for on-beat and off-beat targets, and collapsed over the levels of Predictability.

#### Interaction between beat-based and memory-based expectations

The Predictability and Periodicity of the sequences also affected hit rates in interaction, though depending on the position of the target, as reflected in a significant three-way interaction in the overall ANOVA between Periodicity, Position, and Predictability (Experiment 1: *F*(1,29) = 6.69, *p* = 0.015, η_p_^2^ = 0.19; Experiment 2: *F*(1,30) = 5.59, *p* = 0.025, η_p_^2^ = 0.16) and a significant two-way interaction between Periodicity and Predictability in the separate ANOVA for on-beat targets (Experiment 2: *F*(1,30) = 6.67, *p* = 0.015, η_p_^2^ = 0.18). Off the beat, the simple effect of Periodicity was significant for all comparisons (all uncorrected *p*s < 0.029). Thus, beat-based expectations led to decreased detection of targets when they were presented out of phase with the periodicity and therefore occurred at time points that were mispredicted with regards to the beat. On the beat, the simple effect of Periodicity only reached significance for unpredictable sequences, and only in Experiment 2 (uncorrected *p* = 0.003), showing improved detection through beat-based expectations only in the absence of memory-based expectations. In Experiment 2, we also found an interaction between Periodicity and Predictability for reaction times, with a stronger effect of Predictability in aperiodic than periodic sequences (*F*(1,30) = 6.67, *p* = 0.015, η_p_^2^ = 0.18). The interaction between Periodicity and Predictability did not reach significance in the separate ANOVAs split over position for reaction times (all *p*s > 0.06).

In addition to effects of memory-based and beat-based expectations, in the overall ANOVA we found a main effect of Position for hit rates (Experiment 1: *F*(1,29) = 99.9, *p* < 0.001, η_p_^2^ = 0.78; Experiment 2: *F*(1,30) = 100.3, *p* < 0.001, η_p_^2^ = 0.77) and reaction times (Experiment 1: *F*(1,28) = 37.1, *p* < 0.001, η_p_^2^ = 0.57; Experiment 2: *F*(1,30) = 14.4, *p* = 0.001, η_p_^2^ = 0.32). Targets on the beat were detected more often and faster than targets off the beat, even in aperiodic sequences, most likely reflecting differences in surface grouping.

Thus, behaviorally, memory-based expectations improved target detection both in terms of response accuracy and response speed (see Figure 3). Beat-based expectations similarly lead to improved target detection for targets at expected time points (on-beat), but only when no memory-based expectations were present, and only in terms of response accuracy, not response speed. The improvements in target detection afforded by beat-based expectations were thus small and dependent on memory-based expectations. In contrast, at unexpected time points (off-beat, “mispredicted”), beat-based expectations lead to decrements in performance, both in the absence and presence of memory-based expectations. Interestingly, while we found an interaction between beat-based and memory-based expectations, it was in the opposite direction of what we expected: rather than enhancing each other, when both types of expectations were present, their effects were diminished.

### EEG results

Figure 4 shows the auditory evoked potentials for all conditions. Average amplitudes as extracted from the P1 and N1 time-windows and fronto-central region of interest are depicted in Figure 5.

**Figure 4.**
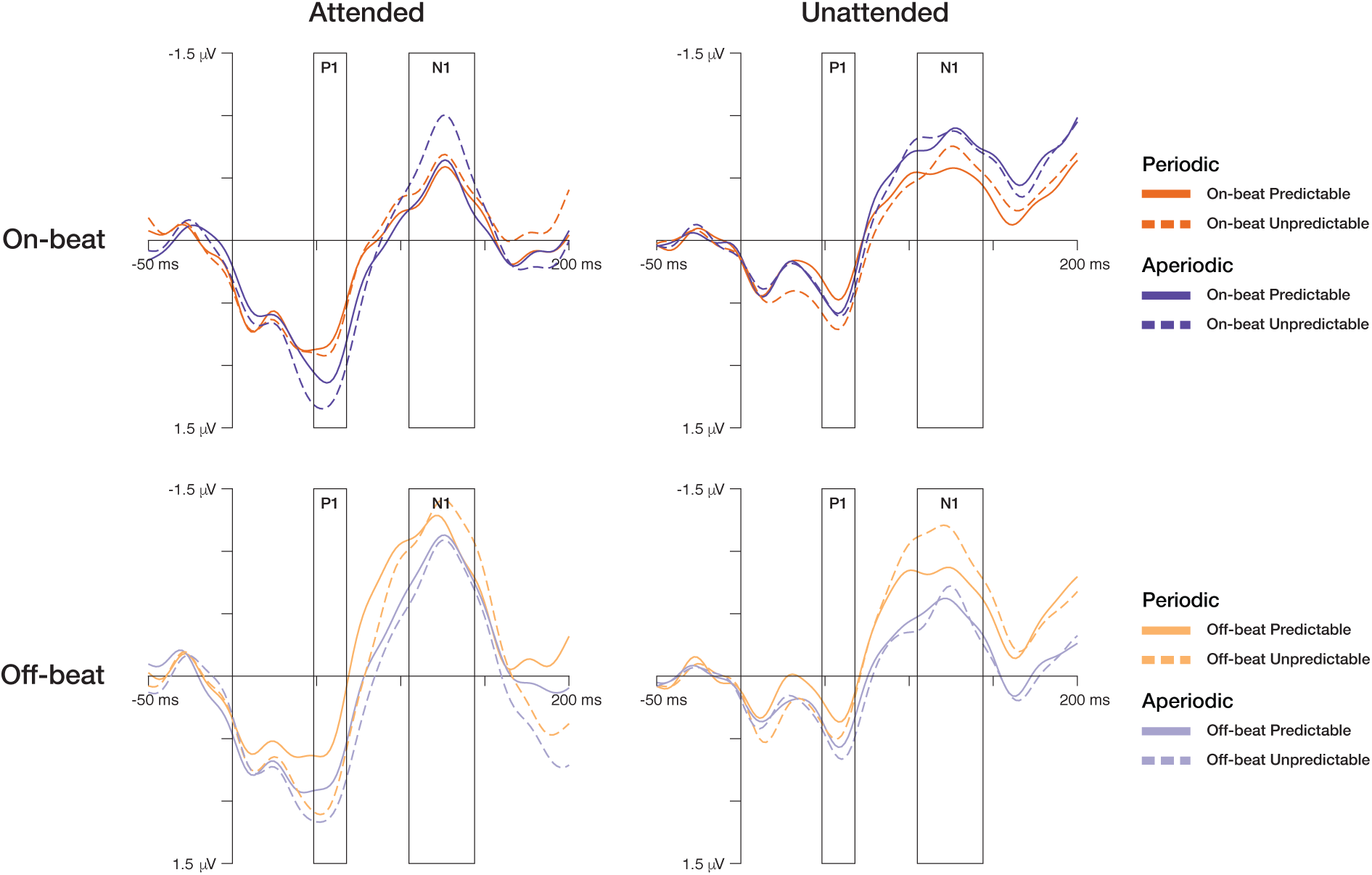
Temporal expectations reduce auditory responses. Temporally expected events evoked smaller auditory P1 and N1 responses, regardless of whether they were attended (task-relevant) or not. Grand-average waveforms are shown for each condition separately. The time windows used for the ANOVA analyses of P1 and N1 are indicated with rectangles. Both memory-based and beat-based expectations attenuated P1 and N1 responses to expected events. N1 responses were additionally enhanced for events off the beat in periodic sequences, i.e., when they were unexpected with regard to the periodicity.

**Figure 5.**
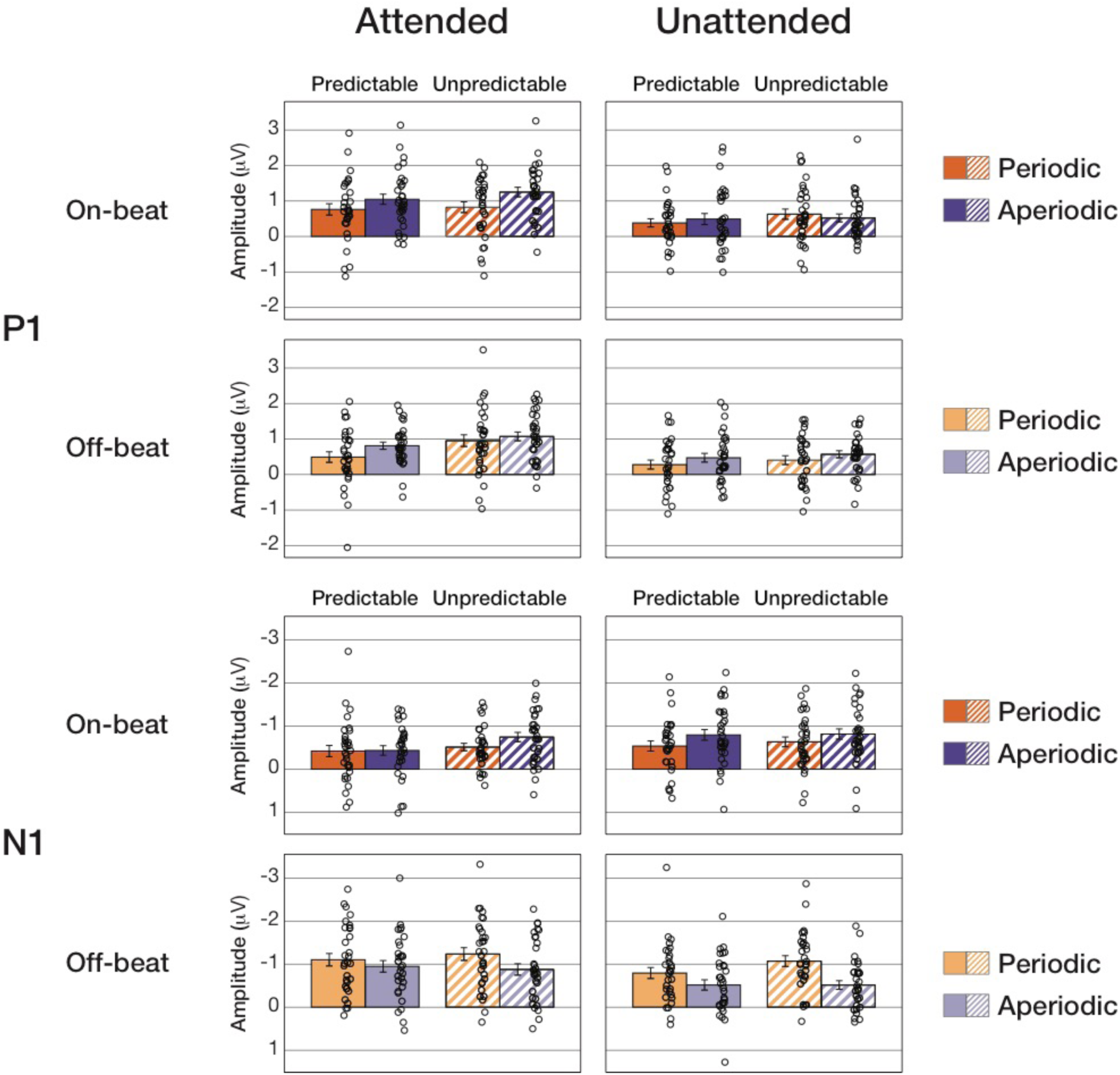
Temporal expectations reduce auditory responses. Amplitudes of P1 and N1 averaged over the time windows used for analysis (see Figure 4) and fronto-central region of interest are shown.

#### Memory-based expectations

As expected, both P1 and N1 responses were smaller for predictable than unpredictable events, reflected in a main effect of Predictability in the overall ANOVA (P1: *F*(1,30) = 10.3, *p* = 0.003, η_p_^2^ = 0.26; N1: *F*(1,30) = 4.52, *p* = 0.042, η_p_^2^ = 0.13), showing attenuation through memory-based expectations. While significant in the overall ANOVA, the attenuating effect of Predictability on the P1 and N1 responses did not reach significance for the separate on-beat ANOVAs (both *p*s > 0.12) and only for the P1 for the off-beat positions (*F*(1,30) = 12.3, *p* = 0.001, η_p_^2^ = 0.29), possibly due to a lack of power when splitting up the data. The overall ANOVA suggested that the effect of Predictability did not depend on task relevance (e.g., none of the interactions including Predictability and Attention reached significance: all *p*s>0.09). However, off-beat, the interaction between Attention and Predictability did reach significance (*F*(1,30) = 4.86, *p* = 0.035, η_p_^2^ = 0.14). Figure 6 shows a summary of the main effects of memory-based expectations on auditory-evoked potentials, collapsed over the different levels of Position and Periodicity. The attenuation of the auditory P1 and N1 responses associated with memory-based expectations suggests processes related to suppression of predicted information, rather than memory-based expectations leading to changes in sensory gain. Numerically, for the P1 response the effects of memory-based expectations were larger in attended than unattended conditions. While statistically the effects of memory-based expectations were independent for task relevance when including all data in one ANOVA, the split ANOVA for off-beat positions suggests that possibly, task relevance does affect the effects of memory-based expectations, at least for the P1.

**Figure 6.**
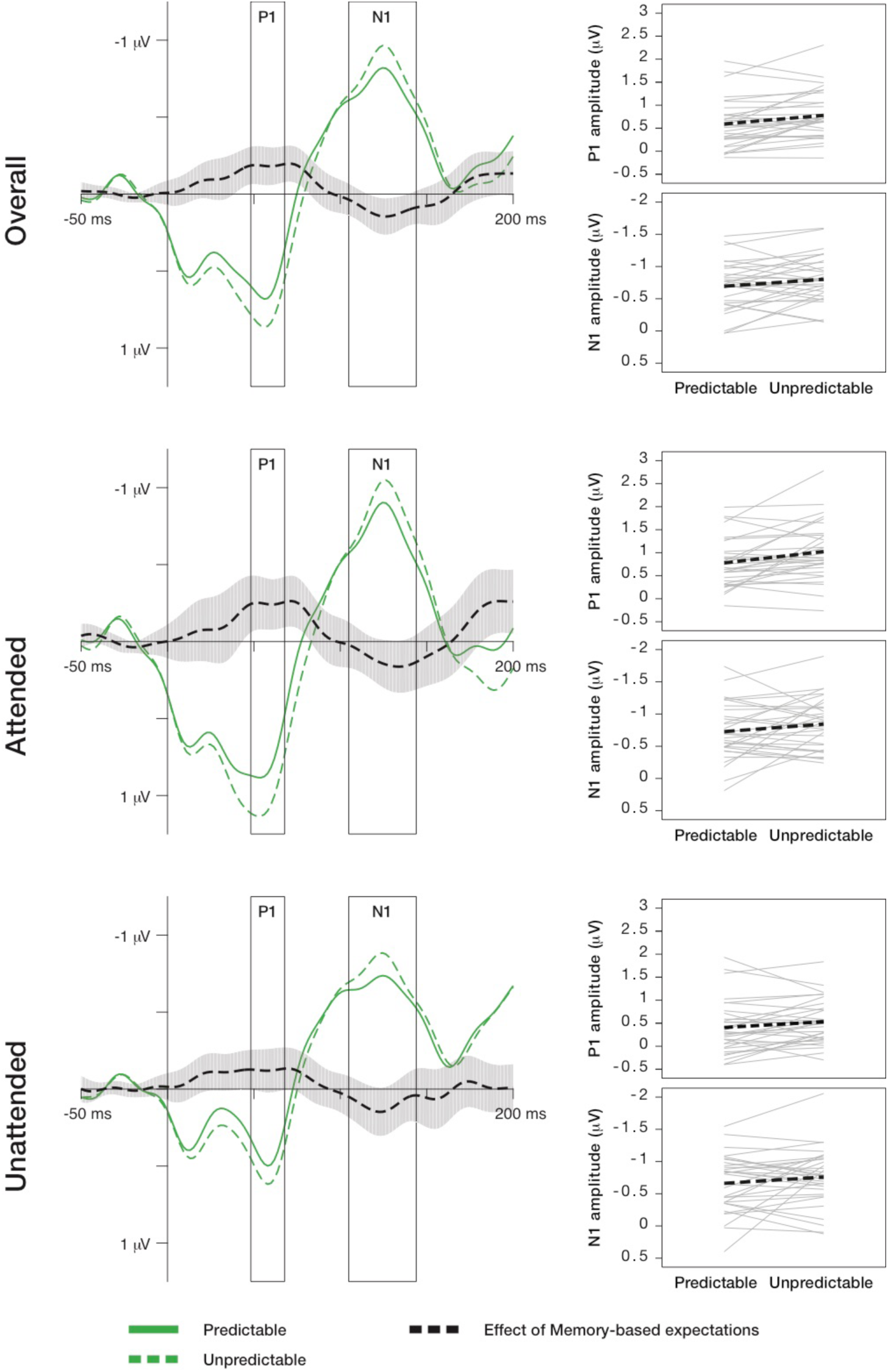
Memory-based expectations attenuated auditory responses. Auditory evoked potentials showing the main effect of Predictability on P1 and N1 responses are shown on the left. The main effect of Predictability is depicted as the difference between responses to events in predictable and unpredictable sequences, collapsed over the levels of Periodicity and Position (i.e., independent of beat-based expectations). In both attended and unattended conditions, P1 and N1 responses were larger for unpredictable than for predictable events. On the right, P1 and N1 amplitudes are shown for all participants separately, with the mean plotted on top.

#### Beat-based expectations

Figures 7 and 8 show a summary of the effects of beat-based expectations on auditory-evoked potentials, collapsed over the different levels of Predictability and separate for responses to events on the beat (Figure 7) and off the beat (Figure 8). Unexpectedly, for P1, the effect of Periodicity did not depend on the position of an event (no interaction between Periodicity and Position: *F*(1,30) = 0.036, *p* = 0.85, η_p_^2^ = 0.001). Instead, we found a main effect of Periodicity (*F*(1,30) = 11.3, *p* = 0.002, η_p_^2^ = 0.27), with smaller P1 responses to events in periodic than aperiodic sequences, significant both on-beat (*F*(1,30) = 5.06, *p* = 0.032, η_p_^2^ = 0.14) and off-beat (*F*(1,30) = 6.72, *p* = 0.015, η_p_^2^ = 0.18). For the N1 response, we did find a significant interaction between Periodicity and Position (*F*(1,30) = 12.7, *p* = 0.001, η_p_^2^ = 0.30). On the beat, N1 responses were smaller to events in periodic than aperiodic sequences (though only marginally so for the N1: *F*(1,30) = 3.99, *p* = 0.055, η_p_^2^ = 0.12). Off the beat, N1 responses were larger to events in periodic than aperiodic sequences (*F*(1,30) = 17.32, *p* < 0.001, η_p_^2^ = 0.37).

**Figure 7.**
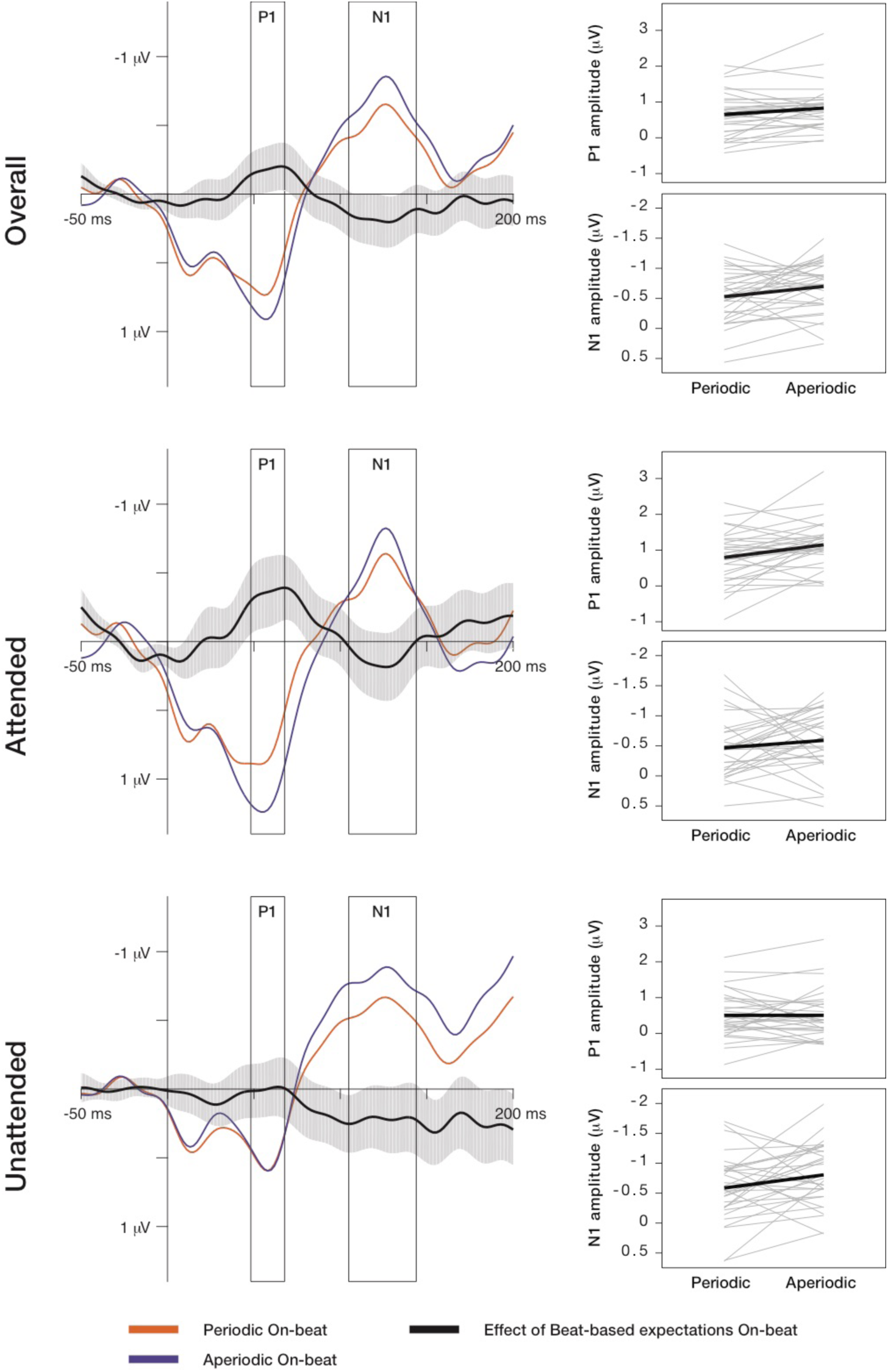
Beat-based expectations attenuated auditory responses (on-beat). Auditory evoked potentials showing the effect of Periodicity on the beat on P1 and N1 responses are shown on the left. The effect of Periodicity is depicted as the difference between responses to on-beat events in periodic and aperiodic sequences, collapsed over the levels of Predictability (i.e., independent of memory-based expectations). In both attended and unattended conditions, P1 and N1 responses were smaller in amplitude for periodic than aperiodic events on the beat. On the right, P1 and N1 amplitudes are shown for all participants separately, with the mean plotted on top. Note that here, the confidence intervals around ERP waveforms represent the variability of the difference waves, in which most of the between-subject variance has been eliminated through subtraction.

**Figure 8.**
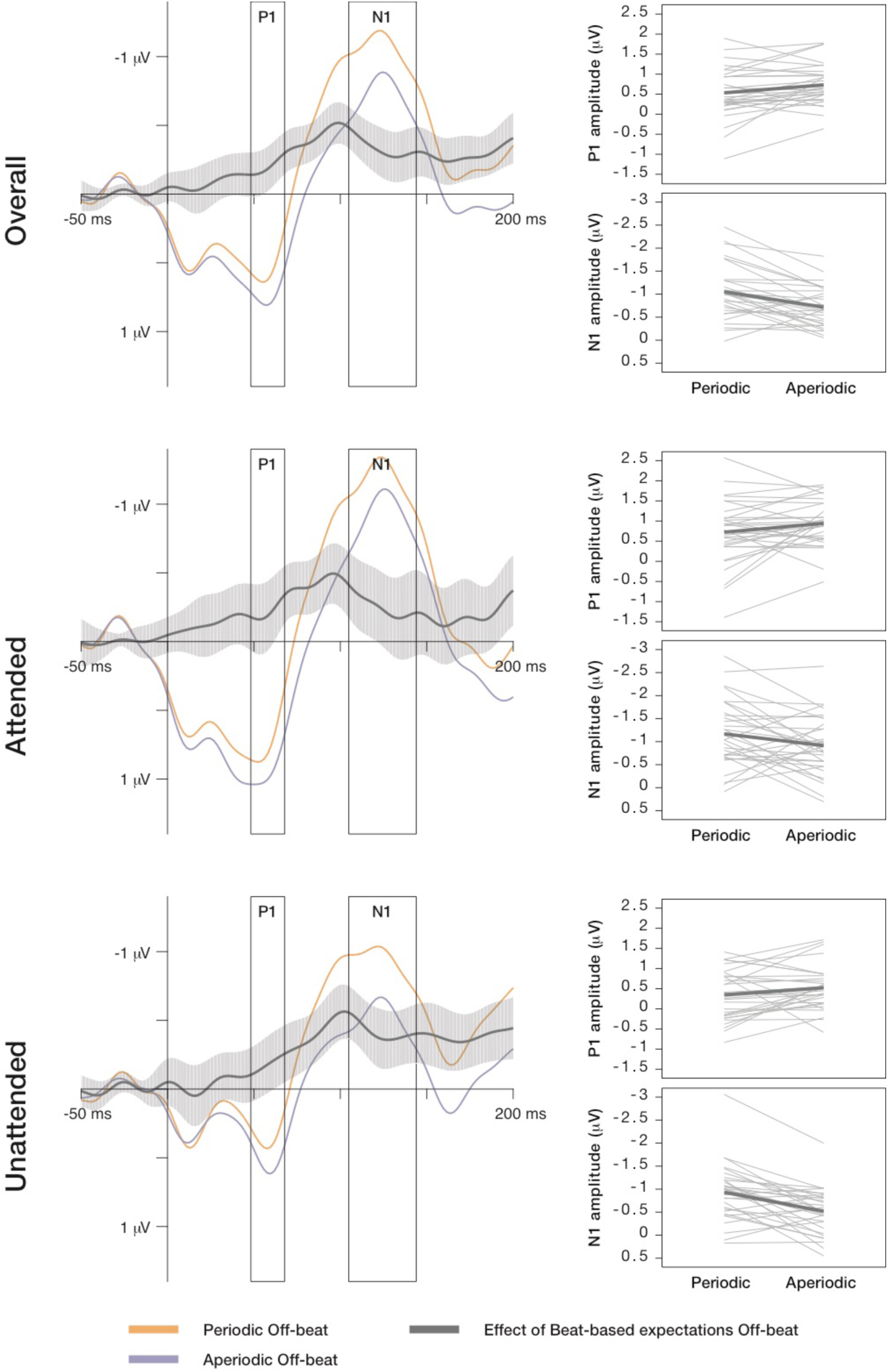
Beat-based expectations enhanced N1 responses for unexpected sounds (off-beat). Auditory evoked potentials showing the effect of Periodicity off the beat on P1 and N1 responses are shown on the left. The effect of Periodicity is depicted as the difference between responses to off-beat events in periodic and aperiodic sequences, collapsed over the levels of Predictability (i.e., independent of memory-based expectations). In both attended and unattended conditions, N1 responses were larger in amplitude for periodic than aperiodic events off the beat, while P1 responses were larger for aperiodic events. On the right, P1 and N1 amplitudes are shown for all participants separately, with the mean plotted on top. Note that here, the confidence intervals around ERP waveforms represent the variability of the difference waves, in which most of the between-subject variance has been eliminated through subtraction.

Like for memory-based expectations, the effects of beat-based expectations were statistically independent of task-relevance in the overall ANOVA (e.g., none of the interactions including Periodicity and Attention reached significance: all *p*s>0.06). Interestingly, similar to the effects of memory-based expectations, while statistically independent of task relevance, the effects of beat-based expectations on the P1 response were numerically driven by the attended condition. Indeed, like for memory-based expectations, when splitting up the ANOVA over the two positions, a significant interaction between Attention and Periodicity was present on the beat for the P1 response (*F*(1,30) = 5.86, *p* = 0.022, η_p_^2^ = 0.16), indicating a small, but significant influence of task-relevance on the attenuation of the P1 response caused by beat-based expectations.

Smaller responses to events in periodic (i.e., beat-based expected) than aperiodic (no beat-based expectations present) sequences, as we found for both the P1 and N1 responses (though the latter did not reach significance), could indicate attenuation of expected events through beat-based expectations, in line with predictive processing, and similar to the effects of memory-based expectations on the P1 and N1 response. In line with this, N1 responses were smallest for events that were expected (i.e., on-beat, in phase with the periodicity), largest for events that were unexpected (i.e., off-beat, “mispredicted”, out of phase with the periodicity), with the amplitude of the N1 to events that were neither expected nor unexpected (i.e., in aperiodic sequences where no beat-based expectations were present) in between. Attenuation of responses to expected events, and enhancement of responses to unexpected events suggests that first order predictions (cf. (Koelsch et al., 2019), rather than changes in sensory gain, underlie the effects of beat-based expectations on perception, similar to the effects of memory-based expectations. However, off the beat, we also found smaller P1 responses to events in periodic than aperiodic sequences, in line with resource withdrawal and smaller responses off the beat in the presence of an underlying periodic beat, as proposed by DAT. This divergent result for P1 responses off the beat is hard to interpret and warrants further research.

In addition to the effects of memory-based and beat-based expectations, P1 responses were larger in amplitude for attended than unattended events (main effect of Attention: *F*(1,30) = 74.3, *p* < 0.001, η_p_^2^ = 0.71, present both on-beat: *F*(1,30) = 39.03, *p* < 0.001, η_p_^2^ = 0.57, and off-beat: *F*(1,30) = 25.85, *p* < 0.001, η_p_^2^ = 0.46). Finally, we found a main effect of Position on N1 responses (*F*(1,30) = 10.3, *p* = 0.003, η_p_^2^ = 0.26). However, the effect of position depended on Attention (interaction between Position and Attention: *F*(1,30) = 15.9, *p* < 0.001, η_p_^2^ = 0.35). In the attended condition, N1 responses to events off the beat were larger than to events on the beat (*p* < 0.001), while in the unattended condition, there was no difference between responses on and off the beat (*p* = 0.74). Likely, these effects of Position, similar to the effects in the behavioral experiments, were due to grouping differences. In line with the interaction between Attention and Position in the overall ANOVA, in the separate ANOVAs for the two levels of Position, the effect of Attention was significant for off-beat positions (*F*(1,30) = 10.0, *p* = 0.004, η_p_^2^ = 0.25), with larger responses in the attended than the unattended condition, and, curiously, for on-beat positions (*F*(1,30) = 6.09, *p* = 0.02, η_p_^2^ = 0.17), for which the responses in the unattended condition were slightly larger than in the attended condition.

To check whether observed results were not confounded by our choice of earlobe reference, we repeated the analysis with mastoid reference, as is more customary in auditory ERPs analyses. Note that these results may have been confounded with the PAMR response, which was present to some degree in several subjects. With mastoid reference, results generally were the same as with earlobe reference. However, with mastoid reference we found a four-way interaction between Attention, Predictability, Periodicity and Position for the P1 (*F*(1,30) = 5.18, *p* = 0.030, η_p_^2^ = 0.15). This four-way interaction did not reach significance for the data referenced to earlobes (*F*(1,30) = 2.64, *p* = 0.11, η_p_^2^ = 0.081). To pursue the interaction in the mastoid-referenced data, we split the data between attended and unattended conditions. In the unattended condition, no significant effects were present. In the attended condition, we found a three-way interaction between Periodicity, Predictability, and Position (*F*(1,30) = 4.50, *p* = 0.04, η_p_^2^ = 0.13). The effect of Periodicity on the P1 response was significant only for events off the beat in predictable sequences (uncorrected *p* = 0.005), and for events on the beat in unpredictable sequences (uncorrected *p* = 0.005). While there were thus small differences in significance between the data sets, the main findings – the effect of Predictability for both P1 and N1 and interaction between Periodicity and Position for N1 – were robust to changes in reference.

#### Comparing beat-based and memory-based expectations directly

Finally, and crucial to our main question, we compared the effects of memory-based and beat-based temporal expectations on auditory-evoked responses directly. We subtracted responses to events on the beat in unpredictable from predictable sequences to index the effects of memory-based expectations, and we subtracted responses to events on the beat in aperiodic from periodic sequences to index the effects of beat-based expectations. Subsequently, we compared the results of this subtraction procedure, both in the restricted time windows around the P1 and N1 peaks, using Bayesian and traditional T-tests, and in a window from 0 to 150 ms after the onset of sound events using cluster-based permutation testing. As the ANOVA yielded somewhat inconclusive results regarding the effects of task relevance, here we report analyses split over attended and unattended conditions, as well as analyses in which we collapsed over attention conditions.

As is visible in Figure 9, the difference waves indexing the two different types of temporal expectations examined have a very similar morphology. Indeed, traditional t-tests showed no significant differences between the effects of Predictability and Periodicity in the P1 window (attended: *t*(30) = 1.24, *p* = 0.23; unattended: *t*(30) = −0.88, *p* = 0.39; combined: *t*(30) = 0.33, *p* = 0.74), nor in the N1 window (attended: *t*(30) = 0.57, *p* = 0.57; unattended: *t*(30) = −0.97, *p* = 0.34; combined: *t*(30) = −0.35, *p =* 0.73). The Bayes factors indicated evidence in favor of the null hypothesis (no difference between conditions), for the P1 window (attended: BF_01_ = 2.6; unattended: BF_01_ = 3.7; combined: BF_01_ = 5.0) and the N1 window (attended: BF_01_ = 4.5; unattended: BF_01_ = 3.4; combined: BF_01_ = 4.9). In other words, the data were 2.6 to 5.0 times more likely to occur under the null hypothesis than in the presence of differences between the effects of Periodicity and Predictability. Bayes factors >3 are regarded as moderate evidence in favor of the null hypothesis (Wagenmakers et al., 2018). The robustness check indicated that the results did not change as a function of the prior used. With the more traditional prior (r = 1), Bayes factors in favor of the null hypothesis ranged from 3.5 to 6.8.

**Figure 9.**
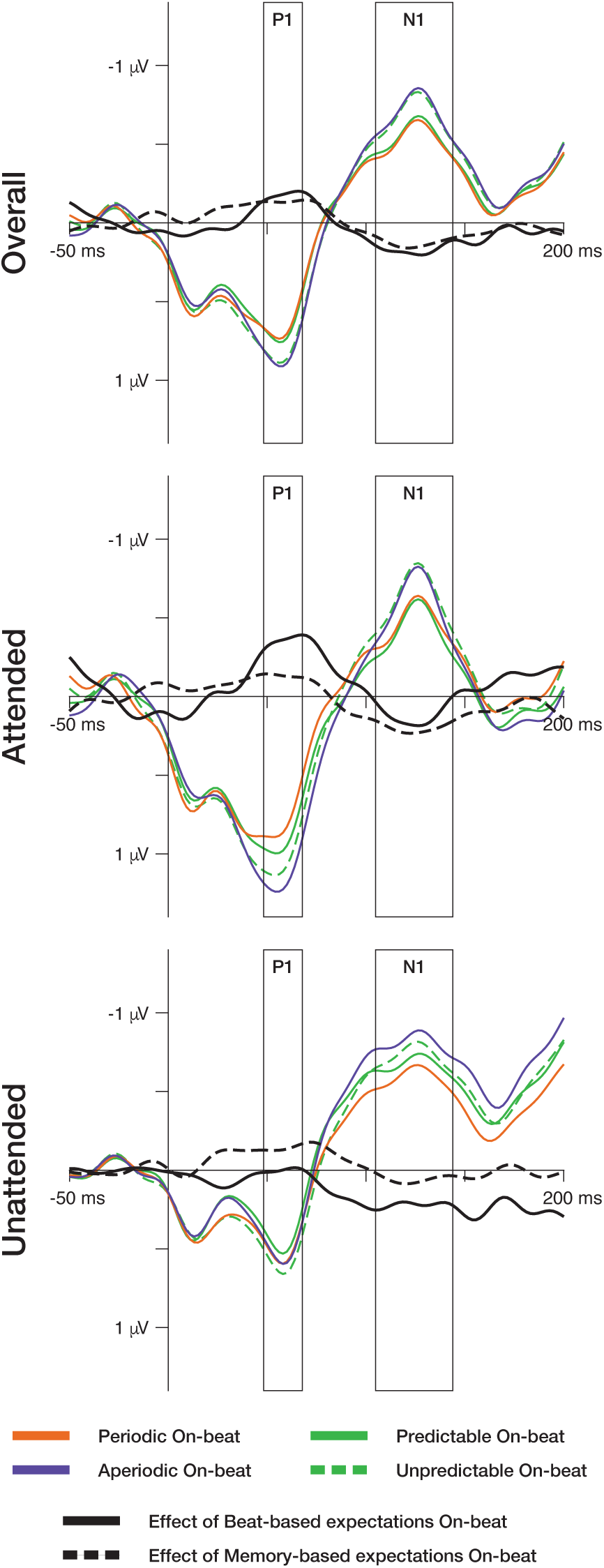
Beat-based and memory-based expectations similarly affected auditory stimulus processing. Effects of beat-based expectations are defined as the difference between responses to events on the beat in periodic and aperiodic sequences. Effects of memory-based expectations are defined as the difference between responses to events on the beat predictable and unpredictable sequences. When comparing the effects of beat-based and memory-based expectations directly using t-tests and cluster-based permutation tests, no significant differences were found.

Cluster-based testing on the ERP waveforms showed no significant differences between the effects of Periodicity and Predictability on the auditory evoked potential, neither in attended, nor in unattended conditions, nor when we collapsed the data over attended and unattended conditions. Thus, using traditional T-tests, Bayesian T-tests, and cluster-based testing, we could not find evidence suggesting differences between the effects of memory-based and beat-based expectations on early auditory responses.

## Discussion

Temporal expectations facilitate sensory processing and perception in dynamic environments, and play an important role in synchronizing our actions to regularities in the outside world, for example, when dancing to a beat in music (Honing & Bouwer, 2019; McGarry et al., 2019). Currently, little is still known about whether shared or separate mechanisms contribute to temporal expectations based on different types of structure in the environment. Specifically, it is unclear whether beat-based expectations (based on periodicity of the input) and memory-based expectations (based on learned predictions of absolute intervals) affect sensory processing and performance in similar ways, independently of each other or in interaction (Breska & Deouell, 2017b; Nobre & Rohenkohl, 2014). Moreover, to what extent these effects rely on task relevance is also still unclear.

In the current study, we show that beat-based and memory-based expectations cannot be differentiated in terms of their effects on auditory processing and/or performance. Both types of expectations lead to enhanced detection of expected events, and to attenuation of auditory responses to those events. Also, beat-based and memory-based expectations interacted, with smaller behavioral effects when both were present. These findings are in line with the notion that beat-based and memory-based expectations are subserved by a shared mechanism for temporal predictive processing (Breska & Deouell, 2017b; Rimmele et al., 2018; Teki et al., 2012). Yet, memory-based expectations enhanced target detection even for events that were fully predictable based on the beat. Also, beat-based expectations lead to reduced target detection and enhanced auditory responses to events out of phase with the periodicity, even when these events were fully predictable based on memory. These findings suggest that while the effects of beat-based and memory-based expectations on sensory processing and performance are the same, the underlying computation may in fact be separate, with beat-based expectations relying (in part) on a rhythmic processing mode characterized by withdrawal of resources off the beat (Breska & Deouell, 2017b). Below, we discuss these findings and their theoretical implications in detail.

Behaviorally, both beat-based and memory-based expectations facilitated target detection in a rhythmic sound sequence, as reflected by enhanced hit rates for targets with expected timing. Although only memory-based expectations improved response speed, the results for beat-based expectations were numerically in the same direction. Thus, unlike previous research (Morillon et al., 2016), we did not find a qualitative difference between beat-based and memory-based expectations on performance. However, in previous work, beat-based expectations were elicited by isochronous sequences, which also elicit memory-based expectations, rendering interpretation of their findings difficult.

That the effects of beat-based and memory-based expectations on auditory processing may rely (in part) on shared mechanisms may further be supported by the observed interaction between the two types of expectations. Beat-based facilitation of detection rates only reached significance in the absence of memory-based expectations. Given that detection rates on average did not exceed 82 percent, even in the fully predictable sequences, it is unlikely that the absence of a beat-based effect here was due to a ceiling effect. Moreover, we expected beat-based expectations to facilitate memory-based expectations, similar to the facilitation of content predictions (“what”) afforded by temporal expectations (Auksztulewicz et al., 2018; Hoch, Tyler, & Tillmann, 2013; Schwartze, Rothermich, Schmidt-Kassow, & Kotz, 2011; Selchenkova, Jones, & Tillmann, 2014). However, if anything, the effect of predictability on target detection was smaller in the periodic than the aperiodic sequences. Thus, when both types of expectations were present, their effects on auditory processing and performance were diminished. This may suggest that beat-based and memory-based expectations also to some extent compete for limited capacity temporal processing to form expectations, leading to interference when both need to be engaged.

In correspondence to our behavioral findings, our ERP results also do not suggest qualitative differences between the effects of beat-based and memory-based expectations. Both P1 and N1 responses to expected events were attenuated, in line with theories about predictive processing, and explained by assuming that the brain only processes input that is not predicted (Friston, 2005; Schröger, Marzecová, et al., 2015). Effects of beat-based expectations are often explained by entrainment models and DAT, which assume increases in sensory gain at expected time points (Henry & Herrmann, 2014; Large & Jones, 1999), leading to enhanced, rather than attenuated sensory responses (Haegens & Zion Golumbic, 2018). Our results, however, do not show enhanced auditory responses due to beat-based expectations.

While the changes in sensory gain suggested by entrainment have often been equated with fluctuations in attention (Henry & Herrmann, 2014; Lakatos et al., 2008; Large, Herrera, & Velasco, 2015), other studies have also shown that entrainment leads to attenuation rather than enhancement of sensory responses (O’Connel et al., 2015; van Atteveldt et al., 2015), like the beat-based expectations in the current study. Moreover, the effects of periodicity, which guides expectations bottom-up, can be dissociated from the effects of task relevance and general top-down attention mechanisms (Kunert & Jongman, 2017). Thus, while entrainment may lead to fluctuations in neural excitability, these may be related to predictions, rather than attention, as proposed by DAT.

Several studies have found enhancement of sensory responses when manipulating temporal expectations, which may seem contradictory to the current findings. However, first, the observed enhancement may depend on the use of different manipulations of temporal expectations. For example, when temporal expectations are cue-based (Hsu et al., 2013), they may indeed be manipulations of the relevance of a time point, rather than prediction (Lange, 2013). Second, when comparing responses to sounds in phase and out of phase with some periodicity, differences in evoked responses may be due to grouping differences between sounds on and off the beat (Schnupp, Rajendran, Harper, Garcia-Lazaro, & Lesica, 2017). Finally, the perception of hierarchical structure in music (meter) may lead to perceived illusory metrical accents (Bouwer, Burgoyne, Odijk, Honing, & Grahn, 2018; Repp, 2010), causing enhanced responses on the beat when compared to off the beat. Indeed, two studies that specifically manipulated beat-based expectations by asking participants to imagine accents on the beat found enhancement of sensory responses (Iversen, Repp, & Patel, 2009; Schaefer, Vlek, & Desain, 2010). To sum up, it may be that entrainment and beat-based expectations are not necessarily related to fluctuations in attention, and that previously reported enhancement of auditory responses is due to grouping and metrical accenting, which we controlled for in the current study by including the aperiodic sequences, and by comparing only responses within one level of grouping. Which specific task- and stimulus-related factors cause auditory responses to be enhanced, and whether attention plays a role in this at all, remains an interesting question for further research.

In addition to considering the effects of beat-based expectations at expected time points, we also looked at how beat-based expectations affected processing at time points that were unexpected, or rather, mispredicted, based on the beat (i.e., off-beat). On these positions N1 responses were larger for events in periodic sequences than for events with similar grouping structure in aperiodic sequences. Enhancement of auditory processing of events off the beat when periodicity is present is in line with these events being more unexpected than their aperiodic counterparts. Behaviorally, we found deteriorated target detection for events off the beat in periodic sequences when compared to aperiodic sequences, in line with reduced processing of events off the beat. Importantly, detection was hampered even if event timing was fully predictable based on learning the sequence of intervals.

Our findings illustrate the importance of also examining effects of beat-based expectations off the beat in distinguishing between beat-based and memory-based expectations. They support the view that beat-based expectations may allow the brain to go into the more efficient “rhythmic mode” of processing, instead of a continuous “vigilance mode” associated with non-periodic input. Rhythmic mode may be accompanied by automatic suppression of out of phase input when entrainment occurs (Schroeder & Lakatos, 2009; Zoefel & Vanrullen, 2017). Beat-based, but not memory-based expectations have indeed been associated with withdrawal of resources from unexpected moments in time, as apparent from immediate CNV resolution after expected time points for beat-based, but not memory-based expectations (Breska & Deouell, 2017b). A rhythmic processing mode characterized by withdrawal of resources off the beat, rather than focusing of resources on the beat, could also explain why beat-based expectations only weakly affected responses on the beat, especially when memory-based expectations were also present. With the current design, we could not probe events that were mispredicted in terms of memory-based expectations. Thus, we cannot be sure that memory-based expectations do not show similar withdrawal of resources from time points where no event is expected. However, focusing on the effects of expectations at unexpected instead of expected time points may be an interesting way to compare beat-based and memory-based expectations in the future.

Off the beat, the enhancement of the N1 through beat-based expectations was independent of task relevance. This may be related to the fact that off the beat, events were mispredicted, rather than unpredicted (e.g., the aperiodic sequences did not allow for the formation of beat-based expectations, making all events unpredicted based on a beat, while in the periodic sequences, events off the beat were not in line with the beat-based expectations that could be formed). The effects of mispredicted stimuli have been shown to be independent of task relevance (Hsu et al., 2018), reminiscent of the independence of the MMN, an ERP response to mispredicted sounds, from task relevance (Näätänen, Paavilainen, Rinne, & Alho, 2007).

For the facilitating effects of both beat-based (on-beat) and memory-based expectations, no interaction with task relevance was present in the overall ANOVA. However, based on the current data, numerically, effects were larger in the attended than unattended condition. Indeed, the interaction with Attention did reach significance for the attenuation of the P1 response when only considering beat-based expectations on the beat and memory-based expectations off the beat, and additionally, for the full ANOVA with mastoid reference. We can therefore not draw strong conclusions about the interaction between task relevance and temporal expectations, other than noting that the absence of an interaction with Attention in the full ANOVA suggests that task relevance does not seem to be a prerequisite for expectations to develop. Whether task relevance interacts with expectations to shape perception is currently a matter of active debate. Recently, several studies have found interactions between prediction and attention, when looking at feature predictions (Auksztulewicz & Friston, 2015; Smout, Tang, Garrido, & Mattingley, 2019), and spatial predictions (Fardo et al., 2017; Marzecová et al., 2017). However, none of these studies looked at temporal predictions, and none found interacting effects as early as the P1 and N1, as studied here. Indeed, a recent study specifically aimed at uncovering earlier effects of attention and prediction found interacting effects of attention and prediction on the P3 in the absence of effects earlier in the processing stream (Alilović, Timmermans, Reteig, van Gaal, & Slagter, 2019). Thus, the presence of an effect of task relevance on expectations may be due to the type of predictions, and the latency of the effect studied, with effects as early as the P1, as found in the current study, possibly only present for temporal expectations. It has been suggested that temporal expectations are fundamentally different from feature and spatial expectations (Auksztulewicz et al., 2018; Sherwell et al., 2017), which would hint that the presence of a possible interaction between temporal expectations and task relevance may indeed be informative of the underlying process. Of note to the current study, while the results with regards to the effects of task relevance were somewhat mixed, they did not in any way differentiate between beat-based and memory-based expectations. Both were to some extent affected by task relevance, and for both, this effect was mainly visible in the P1 response.

In the current study, we strictly controlled the acoustic context preceding the sounds of interest (i.e., the last time point prior to the events used in the analysis that differed in terms of the occurrence of preceding sounds was at −300 ms relative to onset). Yet, we cannot rule out that the longer latency portion of responses to sounds preceding the sounds of interest bled into the ERP waveforms to some extent. In particular, this could have affected the results by affecting the baseline. In addition, if beat-based expectations elicit low frequency entrainment, differences in the phase of the underlying oscillation could lead to a baseline offset. Issues with differences in the baseline can be avoided by using stringent high-pass filtering, eliminating both the beat frequency and the longer-latency portion of the auditory ERP (cf. Chang, Gavin, & Davies, 2012), but such filters have been shown to lead to significant distortions of the P1 waveform, and can even shift low-frequency information from the N1 range into the P1 response (Liljander, Holm, Keski-Säntti, & Partanen, 2016; Tanner, Morgan-Short, & Luck, 2015). While filtering as such is not optimal to eliminate potential confounds in the baseline, here, for the attenuating effects of expectations on both P1 and N1, such a confound is unlikely. Differences in baseline activity would result in a general shift of the ERP waveform, but the attenuation for expected events was present both for P1 and N1 – components of opposite polarity. Thus, our conclusions with regards to the qualitative effects of temporal expectations, attenuation instead of enhancement, are independent of baseline shifts.

For the N1 enhancement we observed for off-beat events in periodic sequences, the effect of baseline differences is harder to rule out, as here, the direction of the effect was opposite for the P1 (attenuation, see Figure 8). This touches upon a fundamental issue when examining temporal expectations in general, and beat-based expectations in particular. Humans are most apt at perceiving a beat in auditory stimuli (Grahn, 2012), at a tempo of around 100-120 beats per minute (London, 2012). When moving away from isochronous stimuli, this will automatically result in rapid successions of sounds, making it difficult to completely eliminate effects of previous sounds in ERPs, even in a paradigm as highly controlled as ours. Also, this prevents the analysis of ERPs after subsequent sounds are presented, in our case 150 ms after the onset of the sounds of interest, even if some effects of expectations may only be visible at longer latencies (Alilović et al., 2019). Balancing the need for ecologically valid stimuli, optimal for eliciting (beat-based) temporal expectations, with the need for strict experimental control allowing for easy assessment of separate ERP components, remains a big challenge for future research (Bouwer et al., 2014, 2016; Honing, Bouwer, & Háden, 2014).

## Conclusion

To sum up, in the current study, we could not differentiate between beat-based and memory-based temporal expectations in terms of their effects on the detection of, and auditory responses to targets with predictable timing, nor in terms of how they were affected by task relevance. Also, beat-based and memory-based expectations to some extent interfered with each other. Similar effects on ERPs and behavior and the presence of interference may point at a shared underlying mechanism, which may be surprising given the evidence from clinical studies (Breska & Ivry, 2018) and research in nonhuman animals (Honing et al., 2018) that suggests that beat-based expectations are distinct from other types of temporal expectations. Indeed, we also found evidence for distinct processing of beat-based and memory-based expectations: When the timing of events was fully predictable based on memory, the presence of beat-based expectations still deteriorated target detection and enhanced sensory responses for events off the beat (at unexpected moments), and when events were fully predictable based on the beat, memory-based expectations still improved target detection.

To reconcile these findings, we propose that beat-based and memory-based expectations overlap in their effect on auditory responses and behavior, in line with both types of expectations serving the same function, but nonetheless, have partly separate underlying mechanisms to form expectations. Future research will have to focus on distinguishing between possible separate underlying mechanisms by examining the neural dynamics and neural networks involved in different types of temporal expectations. This may also elucidate at which point in the processing stream different types of temporal expectations interact, be it at early stages, during the formation of expectations, or at later stages, where expectations exert their effect on perception and behavior. The special status of beat-based expectations, which has been linked with evolutionary advantages of music (Honing et al., 2015), thus remains an open question. We contribute to this question by being the first to look at the orthogonal effects of beat-based and memory-based expectations on responses to auditory rhythm, and by showing how focusing on the effects of expectations on processing at unexpected, rather than expected time points, may provide a fruitful way to differentiate beat-based from other types of expectations in future research.

